# DEADMON: A protocol for long-term monitoring of deadwood-associated species

**DOI:** 10.64898/2026.08.31.747472

**Authors:** J. Purhonen, S. Huttunen, A. Kantelinen, V. Laine, T. Kosonen

## Abstract

Dead wood has a vital role in maintaining biodiversity and ecosystem processes in boreal forests. A large proportion of deadwood-dependent species are threatened by habitat loss and degradation resulting from intensive forest management practices. In Finland, approximately 25% of all forest-dwelling species—around 5,000 species—are dependent on deadwood, of which 596 are red-listed. Currently, there is no large-scale, long-term monitoring programme for saproxylic species in Finland, and the tools required to implement such monitoring remain underdeveloped. Here, we develop, test and report a DNA-based monitoring method for dead wood associated organisms including fungi, lichens, bryophytes, and insects. Three study sites were selected for the research, one from each boreal vegetation zone. From each study site, 16 study logs were selected and permanently marked for long-term monitoring. Our results show that the method is applicable to all studied species groups. However, especially fungi and lichens show considerable difference between species detected morphologically and those detected using DNA-based methods. We conclude that two partly contrasting strategies emerge for implementing long-term monitoring of deadwood -associated boreal species. The first strategy emphasizes as comprehensive geographical sampling of habitat types and tree species as possible. The second approach focuses on long-term monitoring of individual deadwood units across their full decomposition trajectory. We recommend that solutions for unifying the currently fragmented monitoring research on deadwood-associated organisms would be sought together with potential coordinating institutions.

## 1. INTRODUCTION

The significance of deadwood for supporting a wide range of organisms and maintaining ecosystem processes in boreal forests is well established (Jonsell et al. 1998; Stokland et al. 2012; Thorn et al. 2020). A large proportion of deadwood-dependent species are threatened by habitat loss and degradation resulting from intensive forest management practices (e.g. Siitonen 2001; Grove 2002; Figure 1). In Finland, approximately 25% of all forest-dwelling species— around 5,000 species—are dependent on deadwood, of which 596 are red-listed. Moreover, the decline in deadwood availability is considered a threat to approximately one-third (280 species) of threatened forest species (Siitonen 2001; Hyvärinen et al. 2019)

**Figure 1.**
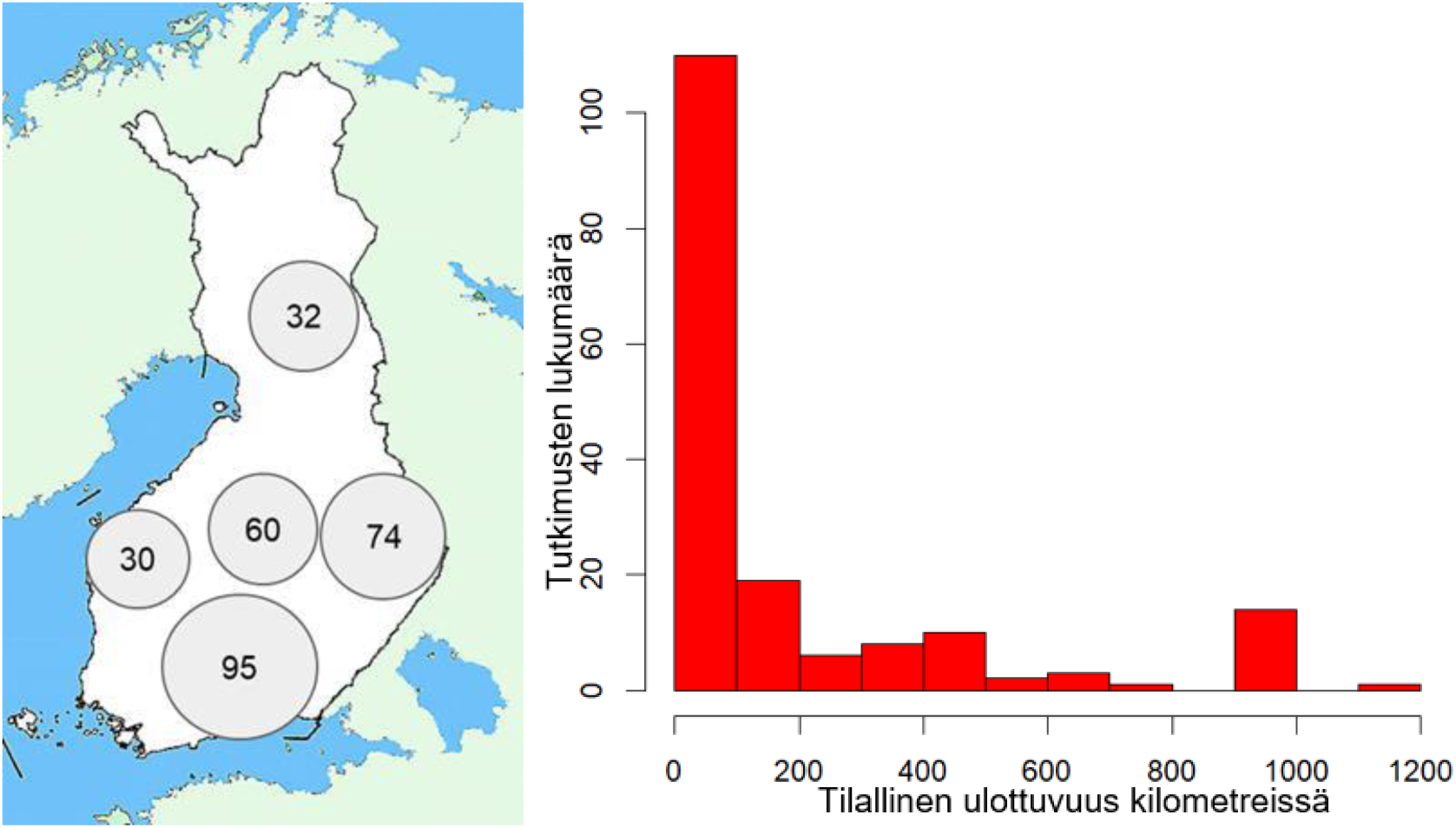
Geographic distribution of studies conducted in Finland (left) and the spatial extent of the study sites (right).

Currently, there is no large-scale, long-term monitoring programme for saproxylic species in Finland, and the tools required to implement such monitoring remain underdeveloped. The need to establish systematic monitoring is further emphasised by the EU Biodiversity Strategy for 2030, which calls for prioritising the identification, mapping, monitoring, and strict protection of all remaining primary and old-growth forests within the EU (European Commission 2021).

This scientific project report describes a DNA-based monitoring method for deadwood-associated organisms, including fungi, lichens, bryophytes, and insects, with particular applicability to boreal and hemiboreal forests.

### 1.1 OBJECTIVE

The main objective of the DEADMON-project is to develop a unified protocol for long-term monitoring of Finland’s deadwood associated species and habitats. In addition, existing methods will be improved and standardized, and new cost-effective tools for monitoring will be tested.

The protocol and survey methods were tested during the summer of 2023 by conducting deadwood associated species inventories (fungi, lichens, mosses, invertebrates) in different types of forest habitats. The structural characteristics of forests and deadwood at the study sites were also surveyed (age/decay stage, tree species distribution, volume, etc.).

The research plan includes preparations for species surveys on approximately one hundred deadwood trunks. Species observations are collected using traditional observation- and specimen-based methods. Species identification uses both morphology-based methods and metabarcoding (mass sequencing) of pooled samples by organism group. In addition, wood dust samples will be collected from the study trunks by drilling. Suitable DNA extraction methods and DNA barcode primers are selected for different sample types (=wood dust, insects, mosses, lichens, fungi).

DNA barcode libraries are expanded to become comprehensive for organism groups especially where the results can be utilized during the project (liverworts, certain fungi, and invertebrates). The applicability and cost-effectiveness of traditional and DNA-based inventory and species identification methods is compared quantitatively and qualitatively in terms of species data coverage, time use, and cost impacts.

#### Expected Results

1. We will define the scientific basis and standardized methods for implementing long-term monitoring.
2. We will determine how deadwood associated species monitoring can be carried out as cost-effectively as possible across space, time, and taxonomic groups.
3. Based on the knowledge produced in the project and existing research data, a handbook of best practices and methods for monitoring will be prepared.
4. We will identify which species are suitable for DNA-based monitoring and which are not, with a particular focus on threatened and indicator species.

### 1.2. DEADWOOD RESEARCH IN FINLAND

To understand the history and presence of deadwood-related research and associated organisms in Finland, we surveyed research using the Dimensions search engine. Studies concerning taxonomy and vertebrates were excluded, as were studies that did not collect or utilize material gathered in Finland. At least one author of each study was required to have a Finnish affiliation. The following search terms were used in the title or abstract: *dead wood*, *deadwood*, *saproxylic*, *epixylic*, *wood-inhabiting*, *coarse woody debris*, and *fine woody debris*. A total of 223 studies were ultimately included. The review will be published as an article.

The topics included in the review article relate to the following themes: basic ecological research, forest management, conservation, decomposition processes, survey methods, population genetics or biology, forest dynamics, and urban ecology. We also examined where and when the research material had been collected, what kinds of habitats were covered, which species were studied, what kinds of data collection methods had been used, what types of deadwood had been investigated and how, and whether DNA methods had been used in the studies.

Most of the studies focused on fungi, arthropods, or deadwood alone (Table 1). Among studies concerning organisms, only a handful included more than one organism group. DNA methods have not yet played a significant role for organism groups other than fungi. Geographically, data collection has been concentrated in southern Finland, while western and northern Finland have received less attention. Most studies cover areas of less than 100 km (Figure 1).

**Table 1.** Number of studies conducted in Finland on deadwood or deadwood-associated organisms in different organism groups, and the number of studies using DNA methods.

| Organism Group | DNA / Total Number of Studies |
| --- | --- |
| Fungi | 17 / 77 |
| Invertebrates | 1 / 64 |
| Mosses | 0 / 12 |
| Lichens | 1 / 16 |
| Deadwood | 0 / 77 |

## 2. MATERIAL & METHODS

### 2.1. STUDY SITES AND SURVEY METHODS

#### Study Sites

Three study sites were selected for the research, one from each boreal vegetation zone:

1. Evo, Kotinen Old-Growth Forest Area (Hämeenlinna, southern boreal vegetation zone),
2. Ulvinsalo Strict Nature Reserve (Kuhmo, middle boreal vegetation zone),
3. Pallas–Yllästunturi National Park (Kolari, northern boreal vegetation zone) (Figure 2).

The sites were selected based on the diversity of deadwood and logistical feasibility, so that the study would include a north–south gradient.

#### Study Logs

From each study site, 16 study logs were permanently marked (Figure 4). The logs were sampled by first randomly selecting a compass direction from the site and then proceeding along that direction. Deadwood was surveyed within an approximately 50-meter-wide belt. The first fallen log meeting the selection criteria was chosen as the study log.

**Figure 4.**
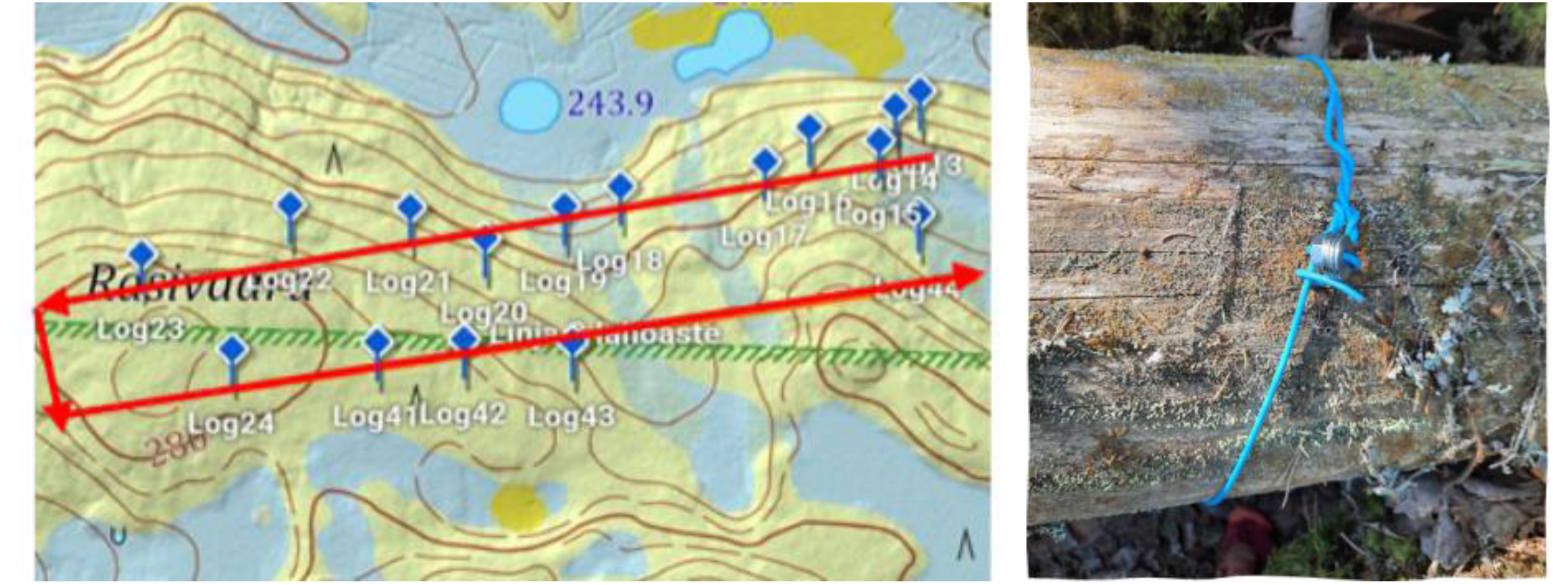
Sampling and marking of study logs using metal tags (photos: Jenna Purhonen).

A minimum distance of 50 meters was required between logs so that, in areas with abundant deadwood, all study logs would not be concentrated in the same location. From each transect, a deadwood log meeting the selection criteria was chosen. The diameter had to be greater than 15 cm, the tree species had to be spruce, pine, birch, or aspen, the minimum log length was 10 meters, at least half of the log had to be lying on the ground, and the decay stage had to be 1, 2, 3, or 4. For each tree species–decay class combination, one log was selected, resulting in a total of 16 logs per site. If no suitable study log was found within 200 meters, the search direction was changed by randomly turning 90 degrees left or right, walking 100 meters, and then returning to the original search direction.

#### Species Inventories

Biological observations were collected from two different parts of each study log: the base and the middle section. In addition, at Pallas–Yllästunturi National Park, a third sampling point (Base 2) was surveyed from logs with decay stages 2 or 4 (a total of 9 logs) (Figure 5).

**Figure 5.**
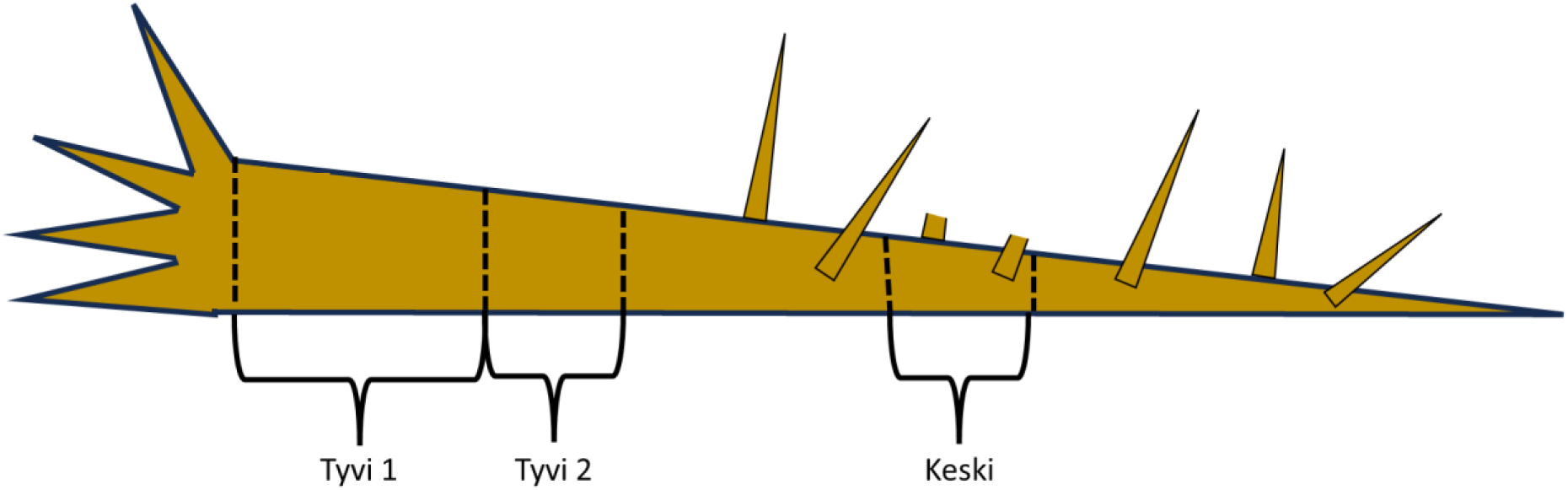
Survey locations on a study log: “Base 1” refers to the first 2 meters from the root collar/break point towards the top of the log; “Base 2” refers to a 1-meter section adjacent to Base 1, further towards the top of the log (only in Pallas–Yllästunturi National Park); and “Middle” refers to a 1-meter section located at the midpoint of the fallen portion of the log, towards the top of the log.

Invertebrates, mosses, lichens, fungi, and slime molds were surveyed from all study logs.

Three types of emergence traps were installed on each study log (Figure 6). At Base 1, a DIY-style trap made of coarse mesh fabric open at the bottom was set up on two sticks. At Base 2, a tent-like bottom-open trap made of fine mesh fabric was installed (MegaView Science Amphibious Emergence Trap BT2008). At the middle section, a tightly fitting trap that wraps around the log was used (Terrapolar Wood Emergence Trap). The traps were left in place for two weeks (from May–June to early July). Insect sampling was conducted on study logs with decay stages 1, 2, and 4.

**Figure 6.**
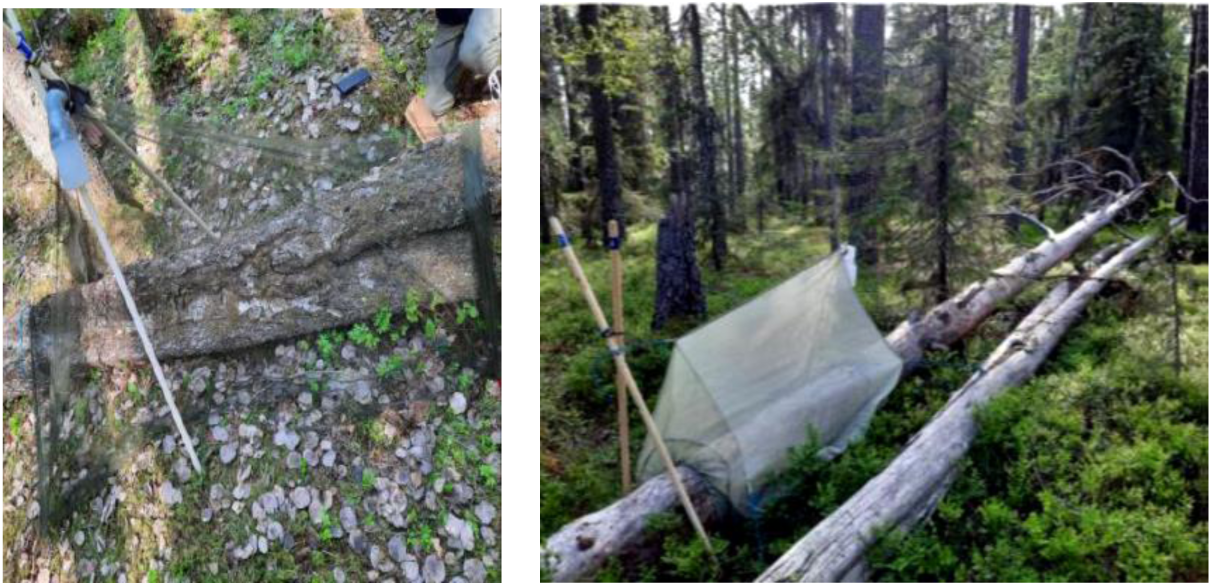
Emergence traps at Base 1 and Middle (photos: Jenna Purhonen).

Surveying the middle section of logs using emergence traps is challenging in tree species that have branches along the entire length of the trunk (such as spruce). The branches must be removed using a saw. In addition, bottom-open traps are difficult to install on logs that are more than half a meter above the ground. Surveying highly decayed logs with emergence traps designed to fit tightly around the trunk is also challenging, as the logs are often already pressed firmly against the ground.

From mid-August onwards, the areas covered by the traps were surveyed for other organism groups (mosses, lichens, fungi, and slime molds). A specimen of every species was collected from each survey location. The samples were dried and used to confirm field identifications later in the laboratory. For each species, a sample was also taken for DNA-based identification using metabarcoding (mixed-species samples), so that all species from a given survey location were included in a single sample. The way these mixed samples were assembled varied depending on the organism group. For mosses, DNA samples can be collected in the field during visual surveys. Accordingly, for mosses, a few shoots per species per survey location were separated in the field into individual sample bags. For lichens, fungi, and slime molds, sampling had to be carried out afterwards in the laboratory using a dissecting microscope. From dried material, a small fragment or a few fruiting bodies (for species with millimeter-sized fruiting bodies) were picked into a single Eppendorf tube per survey location. During the August surveys, wood samples (drill shavings) were also collected from the same locations, particularly to detect fungi that do not produce fruiting bodies.

The insect trapping and other inventories were carried out as follows:

- Evo, Kotinen Old-Growth Forest Area: insect sampling 24.5–9.6.2023; inventory 11.9– 15.9.2023
- Ulvinsalo Strict Nature Reserve: insect sampling 10.6–26.8.2023; inventory 19.8– 23.8.2023
- Pallas–Yllästunturi National Park: insect sampling 27.6–11.7.2023; inventory 24.8– 30.8.2023

#### Laboratory Work

Morphological identification of insect samples was started at Finnish Museum of Natural History (LUOMUS) in September 2023. Entomologist Jaakko Mattila, Master’s student Anna Karellou, and research assistant Maria Reiman reviewed all insect trap samples were reviewed and sorted the insects into broad taxonomic groups and by trap. All beetles were identified to species level, totalling 462 specimens.

A total of 105 samples each were collected for fungi, lichens, and mosses—one from each survey plot. Morphological identification of fungal samples was initiated at the University of Turku by Timo Kosonen and Annika Metso, and at the University of Jyväskylä by Jenna Purhonen and Veera Silenius in October 2023. Annika prepared mixed-species samples for DNA metabarcoding by cutting, scraping (e.g., Corticoid fungi), or picking (e.g., small Ascomycete fungi) a small portion from fruiting bodies into a single sample tube per survey location. Annina Kantelinen identified all collected lichen samples in October 2023 and similarly prepared them as mixed-species samples for DNA-based metabarcoding. Moss identification was initiated by Suvi Leskinen already during fieldwork. In liverworts, oil bodies and other structures were documented under a microscope during identification. Suvi identified liverwort samples together with Xiaolan He and leaf mosses together with Sanna Huttunen. For DNA barcoding, Suvi prepared the samples at the LUOMUS herbarium by collecting a few shoots from each population into separate paper bags per species.

#### DNA Extraction Methods

The choice of DNA extraction method depended on the organism group and sample size. Various commercial extraction kits are available on the market for different sample types. Some can be used directly with standard protocols, while others require minor protocol modifications to ensure successful extraction from specific sample types.

All DNA extractions in the DEADMON project were carried out at the laboratory of Finnish Museum of Natural History (LUOMUS). DNA concentration and purity of the extracted samples were checked using a Nanodrop spectrophotometer. Lichens, mosses, and fungi were extracted using the NucleoSpin™ Plant II kit (Macherey-Nagel), while wood dust samples were processed using the NucleoSpin® Soil kit (Macherey-Nagel). Arthropods were extracted from trap samples using the NucleoSpin® DNA Insect kit (Macherey-Nagel), and individual beetles were processed using the Thermo Scientific™ Phire Tissue Direct PCR Master Mix method, which does not destroy the specimen, allowing beetles to be added to museum collections after extraction.

The extraction protocols were not followed strictly according to manufacturer instructions; instead, minor modifications were made for each sample group. Sample homogenization was performed differently depending on the material: for lichens, fungi, mosses, and wood dust, homogenization was done either manually using a micropestle (lichens) or with metal beads using a TissueLyser device (other groups). For particularly resistant samples, the use of liquid nitrogen at this stage is recommended. For arthropods, the trap material was first sorted into taxonomic groups prior to extraction. Extractions were then performed in groups by combining small-bodied taxa into pooled extractions, medium-sized taxa into separate pools, and for large individuals only a leg or head was used. In cases where a particular insect group was very abundant, it was treated as its own extraction batch. The protocol recommends 40 mg of sample per extraction, and therefore pooling had to be adjusted according to species size and biomass. Groups with very high individual numbers, such as ants and dipterans, were homogenized using an electric coffee grinder (Senz SECGOK2008), and 40 mg of the resulting material was used per extraction. For future applications, if sample biomass is large, it is recommended to use extraction methods that allow larger volumes, such as 20 ml or 50 ml Falcon tubes. However, performing extraction and sequencing at different size classes improves detection of small species within the samples.

From the insect samples, 43 beetles were selected for DNA barcoding. A non-destructive extraction method was used, allowing the specimens to be preserved and later added to the collections of Finnish Museum of Natural History (LUOMUS) after extraction. For mosses, 36 samples were extracted for DNA barcoding using the NucleoSpin Plant II kit (Macherey-Nagel). Half of the insect samples from the Evo site were processed in 2023, and the remaining half in early 2024. Extraction of wood dust samples, as well as moss and lichen samples, was also initiated in November–December 2023. Research assistant Maria Reiman performed the extractions and prepared the samples for sequencing.

#### DNA-based species identification

Deadwood-associated biota can be identified using both morphological and DNA-based methods. DNA barcodes from individual specimens can be sequenced and compared against a reference library, or the DNA barcodes of all organisms living in deadwood can be sequenced simultaneously in a taxon-specific manner and compared to a reference database (metabarcoding), thereby producing a comprehensive species list (Cristescu 2014).

#### DNA barcoding

DNA barcoding refers to a region of DNA in an organism’s genome that is unique to a species and varies as little as possible between individuals of the same species. These regions vary among taxonomic groups (Antil et al. 2022). In the DEADMON project, both mosses and beetles were DNA-barcoded using the Sanger method at the Institute for Molecular Medicine Finland (FIMM (Institute for Molecular Medicine Finland)) using an ABI3730xl DNA Analyzer. Curated sequences will be published in reference databases, including BOLD and GenBank (all BOLD sequences are also available in GenBank). The FinBOL – Finnish Barcode of Life initiative is a Finnish project aiming to produce DNA barcodes for all Finnish species. FinBOL is part of the international International Barcode of Life (iBOL) initiative, which has the long-term goal of building a DNA barcode library for all species globally. DNA barcode sequences produced in the DEADMON project will also be included in the FinBOL database.

In plants and fungi, the internal transcribed spacer (ITS) region of nuclear DNA is commonly used as a barcode region, often divided into ITS1 and ITS2. Chloroplast genes such as rbcL and matK can also be used. In the DEADMON project, the following PCR primers were used to amplify barcode regions in mosses: For the trnL–F region: trnLf (CTCGTGTCACCAGTTCAAAT; Taberlet et al. 1991) and trnC_diplo (5’-CGRAATTGGTAGACGCTACG; Olsson et al. 2009) For rbcLa: rbcLa_F (ATGTCACCACAAACAGAGACTAAAGC) and rbcLa_R (GTAAAATCAAGTCCACCRCG3). For ITS2: ITS_BRY3d (CAACTCTCARCAACGGATA) and ITS_BRY4d (CTTAKTGATATGCTTAAAYTC).

In animals, the mitochondrial cytochrome c oxidase subunit I (COI) gene is commonly used due to its reliability and sufficient resolution. In beetles, the PCR reactions used universal insect primers LCO 1490(F) (GTCAACAAATCATAAAGATATTGG) and HCO2198(R) (AAACTTCAGGGTGACCAAAAAATCA) (Folmer et al. 1994; Sharma et al. 2014). This primer pair works for a very wide range of insect species, including beetles and many other animal groups. It amplifies the first half of the COI gene, approximately 700 base pairs in length. If the DNA sample is highly degraded, sequencing the same region in smaller fragments is recommended, for example using Lep primers (Hajibabaei et al. 2006).

#### Metabarcoding

Metabarcoding also targets sequencing the same barcode regions, but in this approach, sequencing is performed from mixed DNA samples containing multiple species simultaneously. This is a highly efficient method for monitoring biodiversity, although it requires more resources in terms of both funding and bioinformatics expertise (Coissac et al. 2012). In the DEADMON project, metabarcoding (for all organism groups) was performed using DNA from these taxa, as well as environmental DNA extracted from deadwood wood dust. All specimen-derived DNA samples (lichens, mosses, fungi, and arthropods) were sequenced at FIMM (Institute for Molecular Medicine Finland).

A two-step PCR protocol was used, with the first PCR performed at Finnish Museum of Natural History (LUOMUS). This allows verification (e.g., via gel electrophoresis) of which samples successfully amplified before proceeding to the next step. For the first PCR, sequencing tags were added to both primer pairs, to which indexing sequences are attached in the second PCR step. Successful samples were then transferred to FIMM, where the second PCR and sequencing were performed. All sequencing runs were conducted using an Illumina MiSeq platform with 2 × 300 bp reads. The primers used were ITS3 and ITS4 (White et al. 1990) for lichens, mosses, and fungi, and BF3 and BR2 (Elbrecht & Leese 2017) for arthropods. ITS3 and ITS4 amplify the ITS2 region, approximately 350 bp in length. BF3 and BR2 amplify an approximately 460 bp region of the COI gene and represent one of the longer fragments used in arthropod metabarcoding. While longer fragments can improve taxonomic resolution, a large proportion of sequences may be lost during bioinformatics quality filtering because R1 and R2 reads do not overlap in longer fragments; in such cases, only R1 reads can be analyzed. Alternative primer options for insect datasets are presented in Elbrecht et al. (2019).

Environmental samples (wood dust) were sequenced by the company Bioname. DNA samples were sent from Finnish Museum of Natural History (LUOMUS), and Bioname prepared the sequencing libraries and carried out sequencing using an Illumina NovaSeq6000 platform (SP Flowcell, 2 × 250 bp), with each sample run in duplicate. The primers used were fITS7 (GTGARTCATCGAATCTTTG) (Ihrmark et al. 2012) and ITS4 (White et al. 1990).

In metabarcoding, the primary sequencing platform used is the MiSeq system, which was also used for specimen-based sequencing in the DEADMON project. However, when sample numbers increase to several hundred or even thousands, this method becomes relatively slow and expensive. An alternative platform is the high-capacity NovaSeq6000, which was used in the DEADMON project for wood dust samples. Sequencing depth on this platform can be up to 700 times higher, whereas the MiSeq system allows sequencing of longer DNA fragments (300 bp vs. 250 bp), which can improve species-level resolution. Recent comparisons have nevertheless shown that NovaSeq can outperform MiSeq in metabarcoding applications (Singer et al. 2019), and it is possible that future studies will increasingly shift toward using this platform.

All metabarcoding sequence data were analyzed using the QIIME2 software (<u>QIIME2</u>; Bolyen et al. 2019; version 2024.2-amplicon) in the Puhti computing environment of CSC – IT Center for Science. QIIME2 is primarily designed for microbiome sequence analysis, but with minor modifications it is also suitable for other metabarcoding datasets. Both Amplicon Sequence Variant (ASV) and Operational Taxonomic Unit (OTU) clustering approaches were applied to all datasets; however, only ASV-based results were used in the final analyses, as they have been shown to provide a more accurate representation of species diversity (Callahan et al. 2017). In our dataset, both approaches produced very similar results. For taxonomic assignment, the UNITE database was used for ITS data (version 10_dynamic_all_04.04.2024-QIIME2-2024.2), and the COIns database was used for COI data (version 2023.5). When these databases did not provide sufficiently precise species-level identification, BLAST searches were additionally performed against the full GenBank reference database to obtain more accurate assignments. In retrospect, UNITE and COIns are well suited for rapid biodiversity assessments, but for more detailed taxonomic resolution, the use of BLAST is recommended.

## 4. RESULTS

### Arthropods

At Pallas–Yllästunturi National Park, three insect traps failed due to a storm. Overall, a good number of insects and other arthropods were collected from all sites, including species not associated with deadwood. At Evo, Kotinen Old-Growth Forest Area, the total insect biomass was clearly lower compared to the other sites. For the remaining samples, sequencing worked well, and all samples produced a sufficient number of sequences for analysis. Nearly all ASV sequences from the mixed samples (a total of 8,727 ASVs) could be assigned at least to the level of Arthropoda or Insecta; only six ASVs remained completely unidentifiable. Within arthropods, 6,996 ASVs were assigned to order level, and 4,616 to species level. After removing duplicate species assignments (as different ASVs may correspond to the same species) and non-insect taxa, a total of 1,072 insect species were identified to species level, and 1,348 species in total (Table 3). In particular, Diptera (flies) represented a highly species-rich group in the dataset. Final curation of the species list would have required input from multiple insect specialists, especially for Diptera. The most common species at Evo was the fungus gnat *Corynoptera verrucifera*, at Ulvinsalo gall midges (*Cecidomyiidae* spp.), and at Pallas–Ylläs the springtail *Entomobrya nivalis*. Some rarer species were also detected, such as the click beetle *Ampedus suecicus* at Ulvinsalo, and unexpectedly the snail *Columella columella* at Pallas–Ylläs.

**Table 2.**
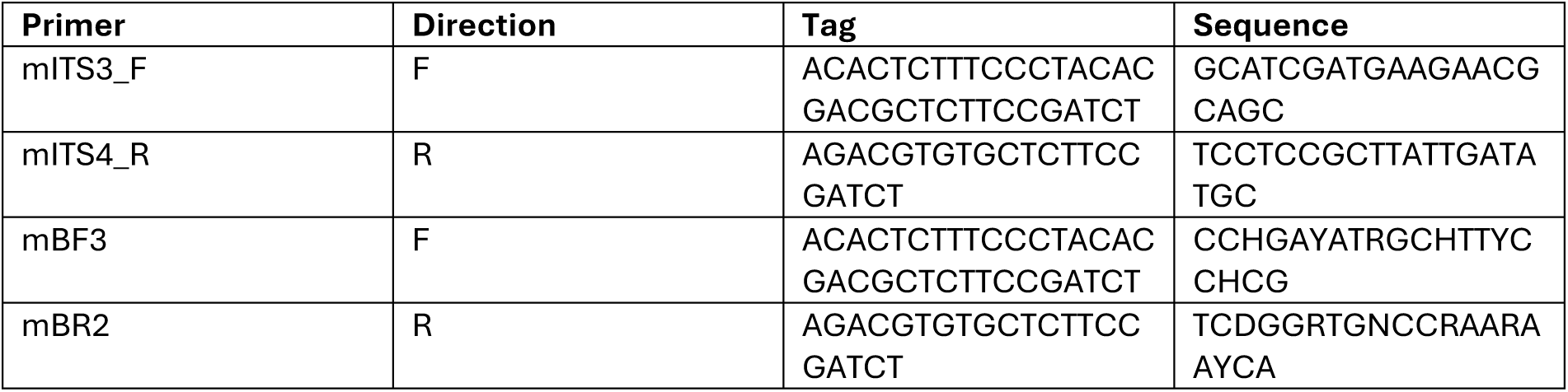
Metabarcoding primers and sequencing tags used (lichens, mosses, fungi, and arthropods).

**Table 3.** Mean species richness per survey location and total species richness across organism groups.

|  | Mean per survey location on trunk |  | Total |  |
| --- | --- | --- | --- | --- |
| Organism group | Morphological identification | DNA Mixture/Wood dust | Morphological identification | DNA Mixture/Wood dust |
| Fungi | 4 | 26 / 75 | ~200 | 770 / 1404 |
| Lichens | 6 | 12 | 65 | 266 |
| Mosses | 4 | 4 | 73 | 54 |
| Arthropods | ~12 | 66 | NA | 1348 |
| Beetles | 7 | 6 | 172 | 158 |

### Beetles

In total, 462 beetle individuals and approximately 170 species were identified based on morphology (Table 3), of which about 130 species are associated with deadwood in some way. A total of 18 traps (K8/U3/Y7) contained no beetles.

From the collected material, 43 beetle individuals were selected for DNA barcoding. Sequences were manually checked and corrected using Geneious software, after which the closest species match was obtained using BLAST against the GenBank database. Most individuals were correctly identified as beetles, but three yielded ambiguous BLAST results. One specimen identified morphologically as the beetle *Otiorhynchus carinatopunctatus* was identified via DNA barcoding as the bark louse *Valenzuela burmeisteri*. This sample was re-extracted using a destructive method, which then produced the correct barcode. The bark louse was likely present as a contaminant in the beetle sample, and because it is softer-bodied than the beetle, its DNA was preferentially extracted using the Phire method. Two other beetle samples were identified via DNA barcoding as belonging to the bacterium Rickettsia, which is also known to occur in insects. Re-extraction did not resolve these cases. In total, 23 high-quality DNA barcodes were submitted to GenBank (accession IDs PQ213101–PQ213123) and later also to FinBOL. At Evo, Kotinen Old-Growth Forest Area, a total of 53 beetle records were obtained, of which 7 were identified only morphologically and not via DNA, and 2 only via DNA. At Ulvinsalo Strict Nature Reserve, beetles were recorded 256 times, with 48 individuals identified only morphologically and 31 detected only through DNA. At Pallas–Yllästunturi National Park, beetles were recorded 138 times, of which 18 were not identified using DNA and 20 were not detected morphologically. Species detected only in DNA-based identification but not in morphological data were often supported by only a few sequences, and are therefore likely misidentifications in some cases. Some of these species are also very small, such as *Scaphisoma agaricinum*, and may therefore be overlooked in morphological sorting of insect bulk samples. Reasons why some species were not detected in metabarcoding but were identified in morphological material may include failed DNA extraction, lack of primer compatibility for that species, or the species not yet being represented in barcode reference databases. In some cases, there were also discrepancies between DNA-based and morphological identifications. Unfortunately, DNA extraction is destructive, meaning that individual specimens cannot be re-examined afterwards to verify identifications. Reference sequence databases also contain errors that are difficult to correct. Metabarcoding-based monitoring data are therefore only as accurate as the reference libraries allow.

### Mosses

Based on morphology, approximately 70 moss and liverwort species were found, of which about one third prefer or require deadwood as a substrate (Table 3). The most species-rich logs were found at Evo, Kotinen Old-Growth Forest Area. Metabarcoding of moss mixed samples yielded 4,690 ASVs, of which 56 were completely unidentifiable. Most of these belonged to fungi (Fungi), totaling 3,587 ASVs. Mosses and liverworts accounted for 370 ASVs, representing 49 different genera or species. Common species in the dataset were well detected by both morphological and DNA-based surveys. However, larger moss species were more readily detected by metabarcoding, while liverworts were poorly recovered. This may be due to factors such as secondary metabolites and their small size. A recommended improvement is to separate small and large mosses and liverworts into different extraction groups. The dataset included a few threatened species, such as *Plagiomnium drummondii* (EN) and *Scapania apiculata* (CR). In addition, metabarcoding detected a species new to Finland, *Lophozia silvicoloides*. For expanding barcode reference libraries, DNA was extracted from 36 moss species and barcoded using three primer pairs. Among these, trnL performed the worst (7 successful PCRs), while rbcLa and ITS performed similarly well (26 and 27 successful PCRs, respectively). Moss DNA barcodes will be published during 2025.

### Lichens

Morphological identification yielded approximately 65 lichen taxa, some of which could only be identified to genus level because they were deficiently developed (e.g. *Cladonia* sp.), and likely include multiple species (Table 3). Metabarcoding of lichen mixed samples performed well. A total of 3,565 ASVs were obtained, of which approximately 1,000 corresponded to lichens or lichen-associated parasites, around 2,200 to other fungi, 340 to plants or algae, and 24 sequences were completely unidentifiable. After curation, 266 genera or species were identified from the dataset. The data included several threatened species (e.g. *Ramboldia cinnabarina, Absconditella celata, Multiclavula mucida*) as well as potentially new records for Finland.

Lichens have evolved multiple times independently, and therefore they occur across several branches of the fungal phylogenetic tree. This somewhat complicates the study of lichens based on DNA data, as they must be separated from other fungi at genus or even species level. According to current knowledge, approximately 100 obligate lignicolous (deadwood-dependent) lichen species occur in the Nordic countries. Facultative species, which can grow on deadwood as well as other substrates, number over 500. Lichen records largely matched expectations, although some common microlichens, such as those in the genus *Micarea*, were unexpectedly rarely detected. This may be due to their inconspicuous appearance in the field.

### Fungi

Based on morphological identification of fruiting bodies, approximately 200 genera or species were recorded (Table 3). The actual taxon richness is likely somewhat higher, as not all fruiting bodies could be identified to species or genus level, for example due to asexual or atypical fruiting structures. In the metabarcoding dataset, 5,464 ASVs were obtained for fungi, of which 68 sequences were completely unidentifiable. Of these, 4,004 ASVs were classified at least to class level within Fungi. In addition, approximately 350 ASVs corresponded to lichens, and 317 ASVs were assigned to Viridiplantae. A total of 34 fungal classes were detected, including 11 classes of Basidiomycota and 13 classes of Ascomycota, representing 24 major classes in total. The groups Basidiobolomycota, Blastocladiomycota, Glomeromycota, Mortierellomycota, Mucoromycota, and Chytridiomycota were excluded from further detailed species-level analyses. A total of 873 unique species or genera were identified, which is four times higher than the number identified morphologically. This difference is partly explained by the fact that metabarcoding detects taxa that do not produce visible fruiting bodies or are present only as mycelium or spores within or on fruiting structures. Threatened species were also detected, such as *Aporpium macroporum* (VU) and *Ionomidotis irregularis* (EN). Slime molds (Myxomycetes) were not detected in the metabarcoding results, as they require a different primer pair targeting the 18S rDNA region (Gøtzsche et al. 2025).

### Wood dust samples

Metabarcoding of wood dust samples differed from the other datasets not only in terms of the sequencing platform but also because each sample was sequenced twice. As a result, the sequencing output was substantially larger than for the other sample groups. When both sequencing runs were combined, a total of 10,779 ASVs were obtained, of which 51 sequences were completely unidentifiable. The majority of the dataset belonged to the fungal kingdom (Fungi), totalling 10,279 ASVs. Of these, 9,031 ASVs corresponded to Ascomycota and Basidiomycota, 483 ASVs to lichens, and 765 ASVs to other fungal lineages. The Ascomycota and Basidiomycota together covered 41 different classes. After curation, approximately 1,400 species or genera of Basidiomycota and Ascomycota were identified (Table 3). From each study site, a single spruce log at decay stage 2 was drilled, and sampling points were taken every 25 cm along the log. A large number of new species accumulated with each additional drilling point, amounting to tens of new species per sample (Figure 7, left).

**Figure 7.**
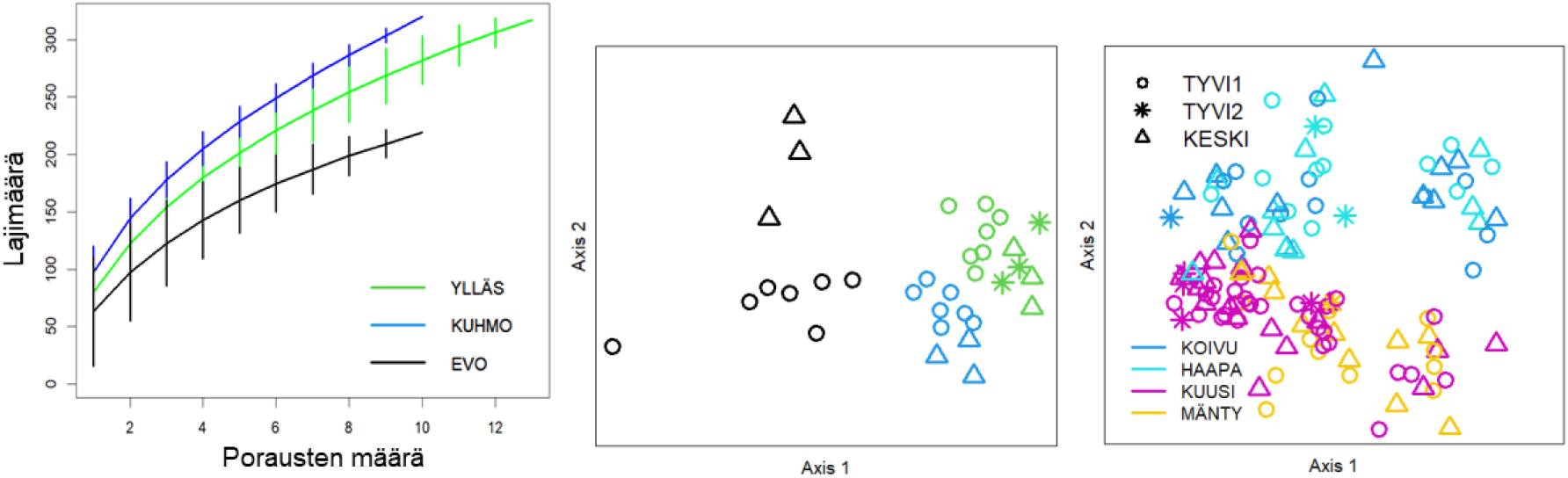
Fungal species accumulation curve with increasing number of drilling points (left), non-metric multidimensional scaling (NMDS) ordination of community composition from frequently drilled logs (middle), and differences in species communities among sampling points in the wood dust dataset (right). Study sites and tree species are indicated by colors, and sampling locations by symbols (Oksanen et al. 2024, R version 4.2.1, R Core Team 2024). In frequently drilled samples, fungal communities differed between logs and sampling points (Figure 7, middle). In the full wood dust dataset, sampling location did not significantly influence fungal community composition (Figure 7, right). This difference is likely due to the fact that the full dataset has not yet been analyzed separately by tree species, which may mask the effects of other factors influencing community composition (koivu=birch, haapa=aspen, kuusi=spruce, mänty=pinus).

### Community composition

Community composition of sampling locations was also analyzed from the mixed-species datasets using non-metric multidimensional scaling (NMDS) ordination (Figure 8). Overall, the results were consistent with those obtained from the morphological dataset.

**Figure 8.**
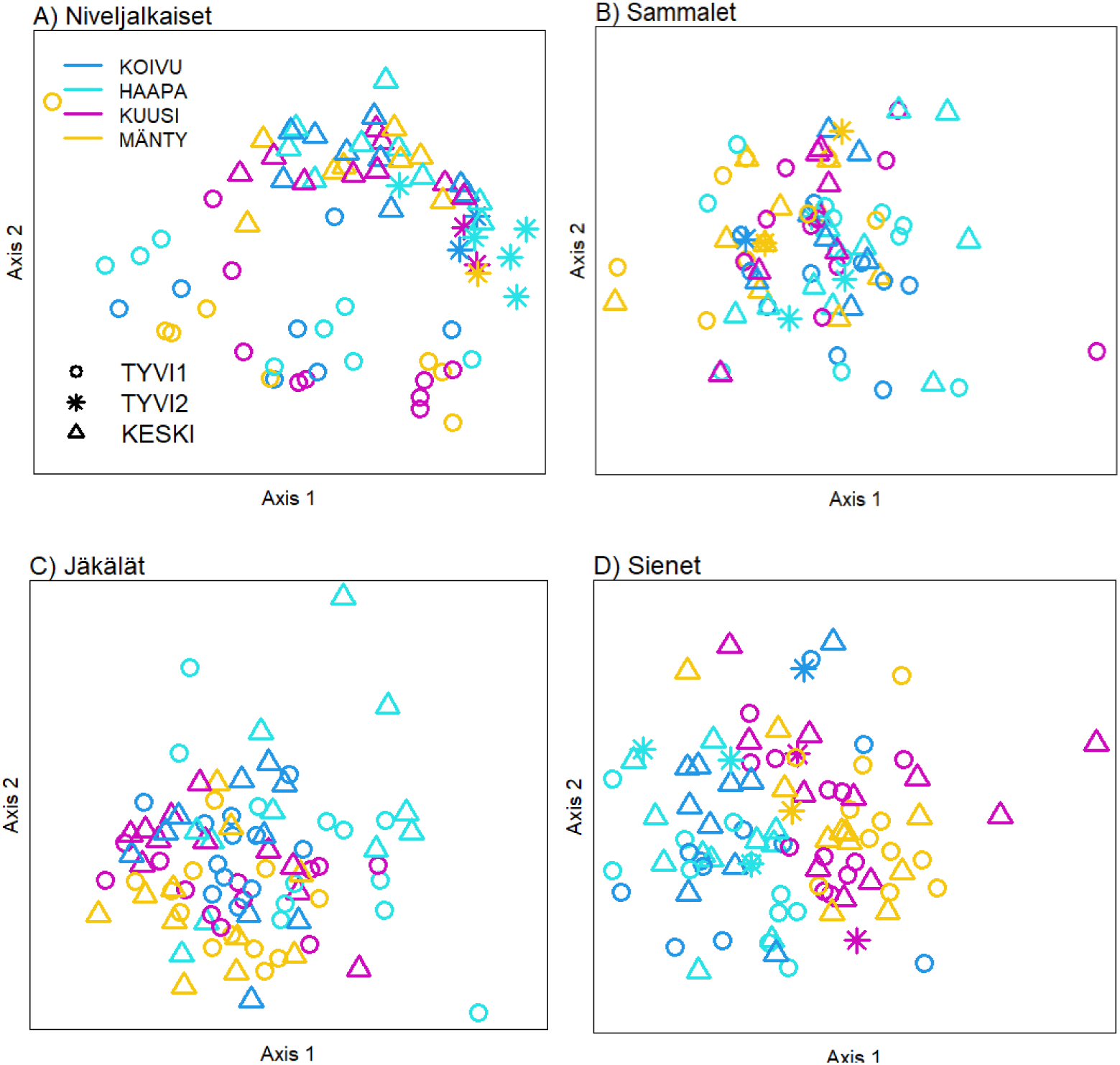
Non-metric multidimensional scaling (NMDS) ordination of species community composition across sampling locations for different organism groups (Organism groups: niveljalkaiset=arthropods, sammalet=mosses, jäkälät=lichens, sienet=fungi. Tree species: koivu=birch, haapa=aspen, kuusi=spruce, mänty=pinus). Tree species are indicated by colors and sampling locations by symbols (Oksanen et al. 2024, R version 4.2.1, R Core Team 2024).

In arthropods, community composition differed clearly between sampling locations. This is most likely due to the use of different types of traps at the sites. For other organism groups, the sampling location had no effect on species composition. Tree species had the strongest effect on fungal community composition, with coniferous and deciduous trees showing more similar species assemblages within their respective groups. Tree species also influenced the community composition of mosses and lichens. In addition, study site had a clear effect on arthropod community composition, and a weaker but still noticeable effect on mosses (Figure 9). For fungi and lichens, the study site did not have a clear effect; however, the datasets still need to be analyzed separately by tree species before this result can be considered conclusive.

**Figure 9.**
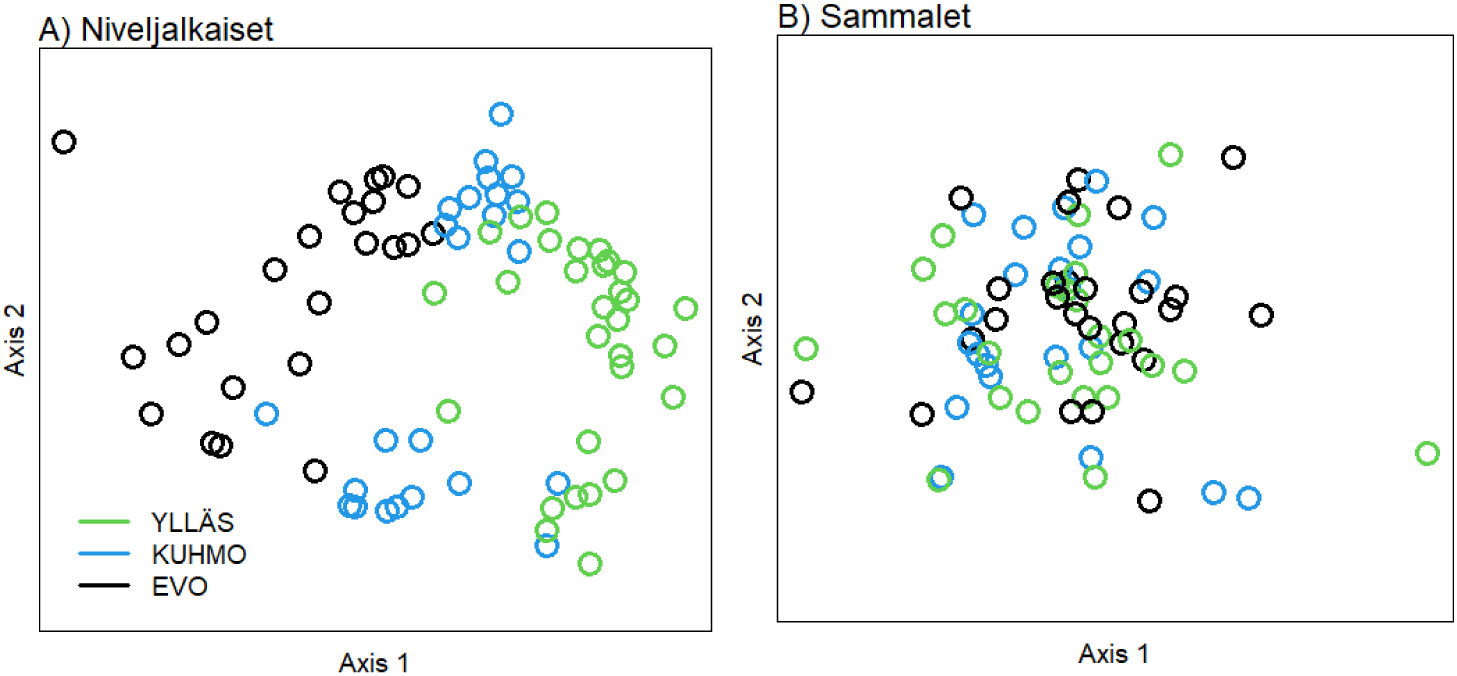
Non-metric multidimensional scaling (NMDS) ordination of species community composition across sampling locations for different organism groups (niveljalkaiset=arthropods, sammalet=mosses). Study site is indicated by color (Oksanen et al. 2024, R version 4.2.1, R Core Team 2024).

## 5. SURVEY AND WORKSHOP

As part of our research, a questionnaire was sent to 45 researchers or experts in Finland working on deadwood and/or deadwood associated organisms. The email survey was distributed together with an invitation to a workshop. It consisted of three questions related to the monitoring of deadwood-associated organisms in Finland. A total of 17 respondents answered the questionnaire. The responses were used as qualitative research material in the project and as support for designing the themes of the workshop.

### Questions

1. Is long-term monitoring of deadwood-associated organisms needed? Please justify: why, or what need it addresses.
2. How should long-term monitoring be implemented? You may, for example, present:
  a. a model where available resources are unlimited,
  b. a model where resources are limited,
  c. your own perspective focusing on a specific organism group.
3. What are the most important research directions, and on the other hand the main challenges, in the study of deadwood-associated organisms over a 5–50 year time horizon?

A one-day workshop was organized at the Botany Unit of the Finnish Museum of Natural History (LUOMUS) in the Nylander Hall on 10 October 2024, from 10:00 to 17:00. A total of 15 participants attended the workshop, including experts representing all organism groups.

### Workshop objectives

1. The primary objective of the workshop was to determine whether there is a need and willingness in Finland to develop and harmonize data collection methods so that datasets collected from different starting points and organism groups would be comparable and consistent.
2. To outline different approaches for implementing long-term monitoring and assess their impact and feasibility.
3. To discuss what deadwood research will look like over a 5–50 year time horizon—who will conduct it, what kinds of data will be collected, and where the data will be stored and used. The workshop also evaluated how comprehensive taxonomic expertise in Finland is and how this expertise can be maintained and secured in the future.

### Workshop summary

- DNA-based methods are constantly evolving, and they suit well for monitoring deadwood-associated biodiversity. DNA barcode reference libraries are constantly improving, and some organism groups can already be detected relatively comprehensively and reliably from environmental DNA (eDNA) data. However, across nearly all organism groups there remains a substantial set of species that cannot yet be detected, most often due to slow progress in foundational taxonomic research.
- Data can be stored as sequence data, but it is also important to consider at what level physical specimens should be preserved and who will be responsible for long-term data management.
- Taxonomic experts remain essential in the future, and DNA literacy is increasingly important for interpreting datasets.
- Prioritising specific habitat types or tree species is challenging, as relevance varies between organism groups.
- It is not advisable to focus only on selected species such as threatened or indicator species, as indicator status can be misleading and we cannot know which currently common species may become threatened in the future.
- Workshop participants were invited to contribute to the writing of a deadwood handbook. The first meeting for preparing the handbook will be held in February 2025.

## 6. CONCLUSIONS AND RECOMMENDATIONS

### Findings and recommendations on inventory techniques

#### Fungi and lichens

For fungi, the difference between species detected and identified morphologically and those detected using DNA-based methods is considerable. The most significant reason is likely that fruiting bodies host a large amount of inconspicuous diversity, including parasites, decomposers, and asexual life stages. In addition, fruiting bodies and deadwood surfaces are subject to substantial spore deposition from both nearby and distant sources. Not all species observed from fruiting bodies were detected in DNA-based mixed samples; on average, one to two species per sample were missed. DNA barcoding-based identification works better for Basidiomycota, typically achieving at least genus-level resolution and often species-level identification. In Ascomycota, identifications typically remained at family level or, at best, genus level. The species richness detected from wood dust samples was remarkably high— substantially higher than that detected from fruiting bodies and/or DNA mixed samples. This difference is likely due to the fact that only a small fraction of the species produces sexual fruiting bodies at the exact time when sampling is carried out. Some of the species observed on deadwood are also such that their fruiting bodies have never been observed at all. In addition, wood dust samples were sequenced at a higher depth, which may also explain the results.

Especially for poorly developed and sterile crustose lichens, DNA-based identification proved useful, as these species cannot always be reliably identified based on morphology alone. Identification of fruticose lichens is also challenging when they are only partially developed. In these cases, DNA-based identification was helpful. Metabarcoding also revealed a visually hidden diversity of fungi and algae that live among lichen thalli.

#### Mosses

Metabarcoding performed best for mosses, and the difference of approximately ten species in total species richness is mainly explained by the fact that liverwort identification from DNA mixed samples was less successful than in the morphological dataset. This could likely be improved simply by separating small and large samples into different tubes (as was done for arthropods). In this way, smaller species—such as many liverworts—would be less likely to be outcompeted during sequencing. Identification and detection of liverworts from mixed samples, or eDNA in general, likely also requires alternative primer pairs, the development of new DNA primers, or the use of multiple primer sets.

#### Arthropods

Among arthropods, only beetles were identified to species level using both morphological and DNA-based methods. The results show that, in addition to mosses, metabarcoding also performed well for arthropods: most of the species identified morphologically were also detected using DNA methods. Of the trapping methods, the Terrapolar wood trap placed in the middle section of the log performed best. The trap is tightly fitted around the log, which minimizes contamination from non-target insects entering from the sides

#### Time requirements of work stages and logistics

DNA-based inventory techniques offer many advantages, but in practice all organism groups require multiple preparatory steps before the actual laboratory phase, and these steps are time-consuming. The appealing idea of preparing samples for DNA extraction already in the field is, in practice, realistic only for mosses.

Field inventories are best carried out in small working groups moving through the study area, which makes it possible to complete inventories for all organism groups—except arthropods—in a single field campaign. A somewhat difficult logistical challenge is that peak activity for arthropods occurs in early summer, whereas for most fungal groups it is concentrated in late summer or even the end of the growing season. Mosses and lichens, in contrast, can be surveyed almost year-round during snow-free periods.

#### Monitoring costs

Table 4 presents an example estimate, based on the project’s experience, of the effort required to inventory 100 logs using the methods applied in this study. The first four columns describe different work stages. The values indicate the average output achievable with one person’s work input per working day. The final column describes the total workload required to inventory 100 logs. In practice, the full workflow—inventorying 100 logs, preparing DNA mixed samples, DNA extraction, and data analysis—corresponds to approximately 1.5 years of full-time work (at minimum, one multidisciplinary field specialist and one highly skilled laboratory technician). In addition, data analysis and curation require further effort, resulting in a total of approximately 300–350 person-days. Material and logistics costs can be estimated at 30–60% on top of personnel costs, along with other indirect expenses. Overall, total costs are on the order of €150,000–€200,000.

**Table 4.** Example calculation of a 100 logs inventory using the methods applied in the DEADMON project.

| **Example** |  |  |  |  |  |
| --- | --- | --- | --- | --- | --- |
| **1pers/trunk/**<br>**day** | **Sample**<br>**collection** | **Morphological**<br>**identification** | **Pre-**<br>**processing** | **DNA**<br>**extraction** | **100 logs**<br>**(excluding**<br>**morphology)** |
| **Emergence**<br>**traps** | 2 | 3 | 5 | 12 | ca. 80<br>working days |
| **Lichens** | 5 | 2 | 3 | 12 | 60 |
| **Fungi** | 5 | 2 | 3 | 12 | 60 |
| **Mosses** | 5 | 2 | 3 | 12 | 60 |
| **Wood dust**<br>**samples** | 5 | - | ? | 12 | 30 |
|  |  |  | **Total 315–340 working days.**<br>The estimate also includes eDNA data analysis, curation, and archiving (5–10 working days per organism group). |  |  |

#### Proposed implementation and recommendations for monitoring

Two partly contrasting strategies emerge for implementing long-term monitoring. The first strategy emphasizes as comprehensive geographical sampling of habitat types and tree species as possible. In practice, this would not allow tracking individual deadwood units throughout their entire lifespan. A geographically comprehensive approach could be implemented, for example, by further developing the Finnish National Forest Inventory (VMI) framework for deadwood. This would involve collecting highly standardized, low-effort, and robust DNA samples from deadwood in a statistically sound sampling design as part of the inventory. If stand-level surveys are conducted using methods such as spore traps or interception traps, it becomes impossible to link detected species to specific deadwood units. As a result, effects of deadwood quality and species interactions cannot be assessed.

The second approach focuses on long-term monitoring of individual deadwood units across their full decomposition trajectory. In practice, this would resemble a fixed research network or a “deadwood laboratory.” Even with a minimal replication design covering main tree species and decay stages, the resulting setup would require several hundred monitored logs. This would limit the number of study areas to a small number (1–5), assuming monitoring is conducted once or twice per decade. Such datasets do not currently exist, and present understanding of decomposition processes and species succession is largely based on chronosequence studies.

There are several types of experimental forests in Finland, mainly within the framework of forestry research, and multiple established research organizations operate in the field of forest science. In contrast, research on deadwood-associated organisms has mostly been based on short-term project funding, resulting in numerous more or less temporary study sites. The field is also relatively fragmented, with researchers and practitioners often working within their own organism-group-specific frameworks, without a coordinating institution or forum. This raises the question of which organization would be capable of coordinating a “deadwood laboratory” and maintaining long-term monitoring over decades. Although the actual field inventory could be carried out by a relatively small number of personnel, the effective use of the resulting data requires a much broader expert network. In this context, it is recommended that all raw data be published openly.

## Acknowledgements

This research was financially supported by the Finnish Ministry of Environment as a part of the research programme BIOMON (2023-2024).

## Notes

### Competing Interest Statement

The authors have declared no competing interest.

